# Decadal trend of plankton community change and habitat shoaling in the Arctic gateway recorded by planktonic foraminifera

**DOI:** 10.1101/2021.08.26.457757

**Authors:** Mattia Greco, Kirstin Werner, Katarzyna Zamelczyk, Tine L. Rasmussen, Michal Kucera

## Abstract

The Fram Strait plays a crucial role in regulating the heat and sea-ice dynamics in the Arctic. In response to the ongoing global warming, the marine biota of this Arctic gateway is experiencing significant changes with increasing advection of Atlantic species. The footprint of this “Atlantification” has been identified in isolated observations across the plankton community, but a systematic, multi-decadal perspective on how regional climate change facilitates the invasion of Atlantic species and affects the ecology of the resident species is lacking. Here we evaluate a series of 51 depth-resolved plankton profiles collected in the Fram Strait during seven surveys between 1985 and 2015, using planktonic foraminifera as a proxy for changes in both the pelagic community composition and species vertical habitat depth. The time series reveals a progressive shift towards more Atlantic species, occurring independently of changes in local environmental conditions. We conclude that this trend is reflecting higher production of the Atlantic species in the “source” region, from where they are advected into the Fram Strait. At the same time, we observe that the ongoing extensive sea-ice export from the Arctic and associated cooling-induced decline in density and habitat shoaling of the subpolar *Turborotalita quinqueloba*, whereas the resident *Neogloboquadrina pachyderma* persists. As a result, the planktonic foraminiferal community and vertical structure in the Fram Strait shifts to a new state, driven by both remote forcing of the Atlantic invaders and local climatic changes acting on the resident species. The strong summer export of Arctic sea ice has so far buffered larger plankton transformation. We predict that if the sea-ice export will decrease, the Arctic gateway will experience rapid restructuring of the pelagic community, even in the absence of further warming. Such a large change in the gateway region will likely propagate into the Arctic proper.

## Introduction

Over the last decades, the Arctic has experienced warming and sea ice decline of “unprecedented” extent, shifting to a climatic state not experienced throughout the 20^th^ Century (Box et al., 2019). A key region for the heat budget and sea-ice dynamics of the Arctic is the Fram Strait. This narrow passage constitutes the only deep-water connection with the Atlantic Ocean, facilitating the inflow of a large portion of the warm and saline Atlantic Water, the export of sea ice from the Arctic (Beszczynska-Möller et al., 2012), and exchange of marine biota between the polar Arctic Mediterranean and the subarctic North Atlantic (Bluhm et al., 2015; Kosobokova & Hirche, 2009; Wassmann et al., 2015). The Atlantic Water (AW) is transported through the eastern part of the strait by the West Spitsbergen Current (WSC), while in the western part, the East Greenland Current (EGC) carries polar water and sea ice from the Arctic Ocean and Nordic Seas to the south (Fig.1). The AW inflow to the Arctic has warmed over the last decades (Beszczynska-Möller et al., 2012; Polyakov et al., 2012; Wassmann et al., 2015) and is regarded as one of the main drivers of the current changes in the Arctic marine environment (Onarheim et al., 2014).

**Figure 1.**
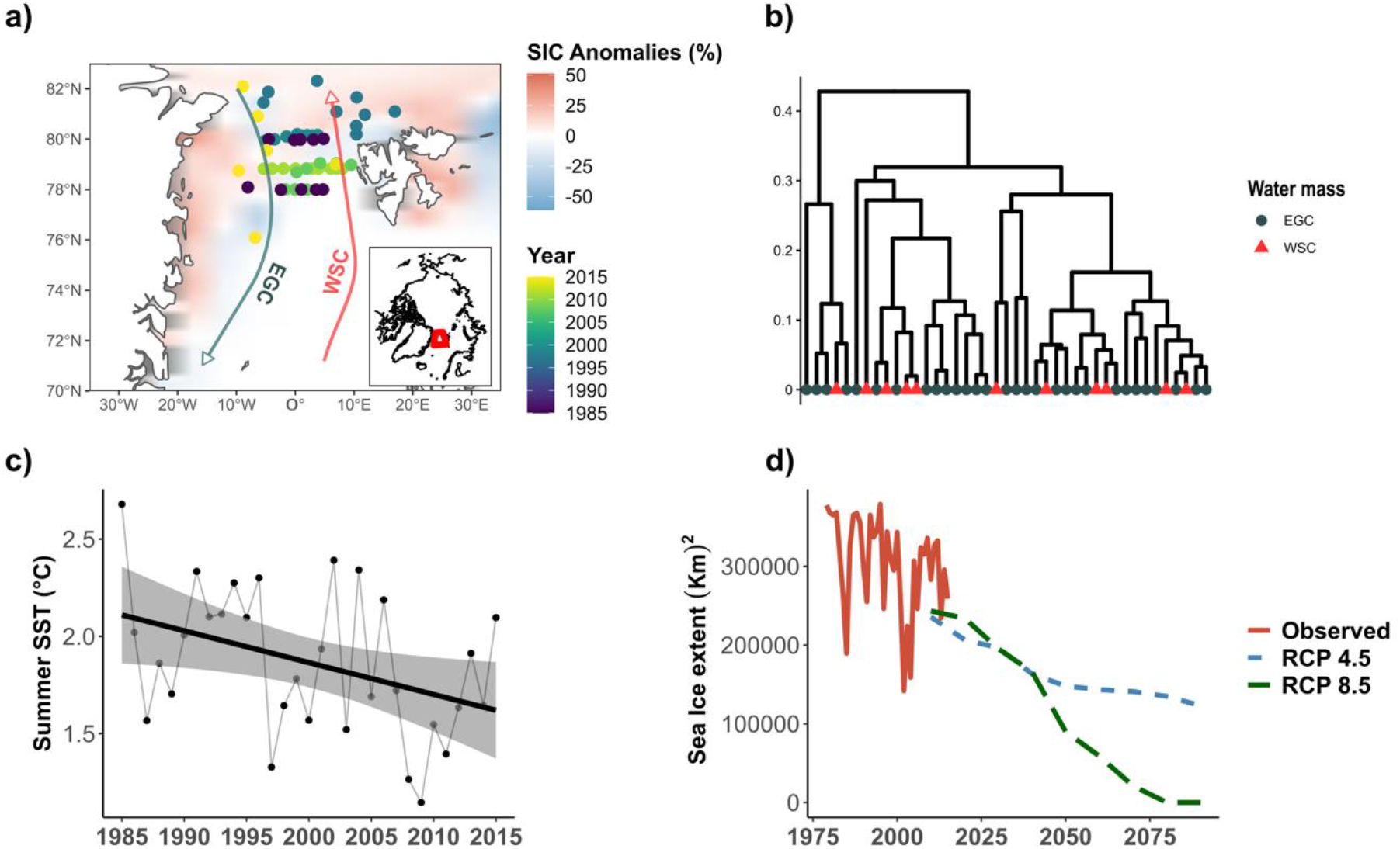
a) Plankton net stations with vertically resolved planktonic foraminifera counts used in this study color-coded by year of sampling. Background colour indicates Sea Ice Anomalies (SIC) in the Fram Strait calculated for the period 1985-2015. Data from Sea Ice Index Version 3.0 (Fetterer et al., 2017). Arrows indicate the two main water masses present in the Fram Strait (East Greenland Current -EGC-and the West Spitzbergen Current -WSC-). b) Hierarchical cluster analysis showing the similarity of foraminiferal assemblages, symbols show the water mass identified with the position of the station. c) Summer SST in the sampling area in the period 1985-2015. Data from NOAA Optimum Interpolation Sea Surface Temperature V2 [weekly resolution] (Reynolds et al., 2002). d) Observational and predicted Sea-ice extent in the sampling area for the month of August from 1979 to 2090. Observational data (red line) from Sea Ice Index Version 3.0 (Fetterer et al., 2017). Model prediction for Representative concentration pathways 4.5 and 8.5 (green and blue lines respectively) based on results of Khon et al. (2017).

The apparent increase in the advection of AW and the resulting changes in the Arctic Ocean are reflected in the “Atlantification” of the marine community in the Fram Strait (Andrews et al., 2018; Gluchowska et al., 2016; Kraft et al., 2013; Schröter et al., 2019). A long-term record of planktonic foraminiferal shells in a marine sediment core from the Fram Strait indicates that the recent changes are unparalleled over the last two millennia (Spielhagen et al., 2011). The ongoing Atlantification of the Arctic gateway contrasts with the changes in the physical environment of the upper ocean in the region. Unlike the rest of the Arctic realm, in the summers between 1985 and 2015, the Fram Strait has been cooling at the surface (~ 0.5 °C), and sea ice has expanded along the east coast of Greenland and Svalbard (Fig. 1). This seemingly counterintuitive trend is a reflection of the increasing reduction in summer sea ice in the Arctic and the associated increased export of Arctic sea ice into the Greenland Sea (Wang et al., 2019). Net changes in sea surface temperatures (SST) in the Fram Strait are therefore the result of the combined effect of increasing advection and warming of northward-flowing AW and increased sea-ice export in the EGC flowing southward. The higher export of Arctic sea ice and its melting in the Greenland Sea also contribute to a large-scale surface freshening, which suppresses oceanic mixing and facilitates cooling of the surface waters (Kwok et al., 2005) that are overlying the warm Atlantic inflow in the subsurface.

Thus, the environmental conditions in the Fram Strait, taken alone, should not facilitate Atlantification of the marine biota. Indeed, the observed increase in abundance of subpolar species and associated community changes have been interpreted as a consequence of warming in the North Atlantic “source” region and intensification of the AW inflow carrying the subpolar biota into the Fram Strait (Wassmann et al., 2015). In this scenario, the increasing proportion of Atlantic biota should occur independently of the local conditions in the Fram Strait and the Atlantification process should be associated with a re-arrangement of the vertical structure of the pelagic communities.

Accurate investigation of these dynamics are missing due to the lack of long-term time series on plankton distribution patterns available from the region (Dornelas et al., 2018). Furthermore, research on zooplankton response to climate change, and foraminifera in particular, rarely include an assessment of the effects on the vertical distribution of the marine species community (Jonkers et al., 2021; Jorda et al., 2020).Here, we analyse three decades of changes in population structure and vertical distribution of planktonic foraminifera, a distinctive group of Arctic unicellular calcareous zooplankton, in the Fram Strait recorded by 51 species-resolved vertical profiles of standing stocks sampled between 1985 and 2015. Planktonic foraminifera species distribution is controlled by temperature (Bé & Tolderlund, 1971; Fenton et al., 2016; Morey et al., 2005), and they show distinct depth habitats, which vary with changing environmental conditions (Greco et al., 2019; Rebotim et al., 2017) and the sedimentary record indicates that foraminifera are sensitive indicators of climate change since the preindustrial era (Spielhagen et al., 2011; Jonkers et al., 2019), making them ideal sentinels of Fram Strait Atlantification and changes in vertical habitat structure. To this end, we combined data from repeated foraminiferal surveys in the Fram Strait with in-situ and regional environmental descriptors to assess the extent and environmental determinants of recent changes in (i) planktonic foraminiferal community composition, (ii) species standing stocks and (iii) shifts in vertical distribution of species.

## Material and Methods

### Biological data

Over the last four decades, the plankton community of the Fram Strait has been sampled regularly with replicate vertical profiles available for most sampling years. Among the collected zooplankton, planktonic foraminifera have been the most frequently quantified and reported at species level, allowing us to compile a dataset of five surveys of planktonic foraminifera repeated at virtually the same location and same time of year in the Fram Strait between 1985 and 2011 containing a total of 45 vertical profiles. In order to extend the length of the time series, we generated new data from one profile taken in July 2014 and a survey with five profiles sampled in July 2015. In 2014, planktonic foraminifera were sampled from three different depth intervals in the upper water column (0–50 m, 50–200 m, and 200– 600 m) by the means of a WP2 net with aperture 0.25 m^2^ and mesh size of 90 μm during an oceanographic cruise with *R/V Helmer Hanssen*. The following year, sampling was carried out on the *R/V Polarstern* using a multiple closing plankton net (Hydro-Bios, Kiel) with an opening of 0.25 m^2^ and equipped with 5 nets each with a mesh size of 55 μm.

Samples from both expeditions were wet-sieved through 250 and 63 μm sieves and stained with Rose Bengal/ethanol mixture after collection to facilitate the distinction between cytoplasm-bearing and empty shells. The samples were processed at the University of Bremen and at UiT the Arctic University of Norway in Tromsø, where planktonic foraminifera were picked under a binocular microscope and air-dried. All specimens in the fraction above 63 μm were counted and identified to species level following the taxonomy of Brummer and Kroon (1988) and Hemleben (1989). Concentrations of the resident species (*Neogloboquadrina pachyderma* and *Turborotalita quinqueloba*) and of the Atlantic species (*Globigerina glutinata*, *Globigerina bulloides*, *Neogloboquadrina incompta*, *Globigerinita uvula*, and *Orcadia ridelii*) were derived from counts by using the volume of filtered water determined from the product of towed interval height and the net opening.

The new data and the literature data had to be first harmonised to the same taxonomy. As a result, counts of *N. pachyderma* and the Atlantic species from the ARK III/3 cruise could not be used in the analyses due to the different taxonomical resolution of the original study (Carstens et al., 1997). In their paper, the authors did not distinguish between *N. pachyderma* and *N. incompta*, previously considered ecophenotypic variants of the same species, but now known to be genetically distinct forms (Darling et al., 2006). Only data on the polar species *T. quinqueloba* collected during the same expedition could be included in the analyses. Because of their consistently low density and variable species composition, the concentrations of all non-resident (Atlantic) species were lumped into one category for the downstream analyses. For samples collected in 2008, the proportion of the Atlantic species was assumed to be 2% of the total assemblage as stated by the authors of the original study (Manno & Pavlov, 2014). Because of the taxonomic lumping, the vertical habitat of the Atlantic species could not be evaluated. Since the distinction between cytoplasm-bearing and empty shells has not been done consistently, the analysis is based on the concentration of all shells. Greco et al. (2019) have shown that this treatment causes a slight but consistent overestimation of the vertical habitat depth, but since the vast majority of the collected specimens in the plankton are cytoplasm-bearing, the effect on standing stock estimates is likely negligible. Further, the different surveys have used different vertical sampling schemes and resolutions. Therefore, the individual vertical density profiles were converted to a common vertical scheme resolving standing stock at three depths (0–50 m, 50–100 m, 100–200 m) using a custom script in R (R Core Team, 2017). This scheme was chosen to avoid extrapolation and reflects the most shared position of depth-interval boundaries among the sampling schemes. We derived total species abundance as the sum of the concentrations within the different intervals. For the two polar species, *N. pachyderma* and *T. quinqueloba*, the depth habitat was calculated as in Greco et al. (2019).

### Environmental parameters

The habitat of planktonic foraminifera reflects the vertical structure of physical and biological properties of the surface ocean layer. Therefore, next to the consideration of the temporal trends, to understand why population densities, species composition and vertical habitat have been shifting, we have tested models explaining the observed variability with physical properties of the environment. In Arctic polar waters, the main parameter affecting planktonic foraminiferal species composition appears to be temperature (Jonkers et al., 2019; Morey et al., 2005) in combination with sea-ice concentration (Carstens et al., 1997; Pados & Spielhagen, 2014). Additionally, in the Fram Strait an important parameter is also the depth of the Atlantic layer (Pados et al., 2015; Simstich et al., 2003). In contrast, salinity, within the range of typical open marine conditions, has been shown not to affect planktonic foraminifera (Greco et al., 2019).

In-situ temperature profiles were retrieved from CTD data from the respective expeditions. Data deposited in PANGAEA were accessed using the R package “pangaear” (version 08.2) (Simpson & Chamberlain, 2018). For nine stations from the iAOOS and HH14 cruises, CTD data were obtained from the original investigators. The CTD temperature profiles were used to extract Sea Surface Temperature (SST), here defined as the average temperature in the uppermost 6 meters from the sea surface, and the minimum depth of the Atlantic Water layer (AWz), defined as the depth where temperature rises above 2 °C (Beszczynska-Möller et al., 2012). As no CTD data were collected during the ARK III/3 cruise (Carstens et al., 1997), for these stations we extracted the SST and AWz from the NOAA Optimum Interpolation Sea Surface Temperature V2 [weekly resolution] (Reynolds et al., 2002) and the Hadley Centre EN4 dataset (Good et al., 2013), respectively. For all stations, in situ sea-ice concentration and the distance from the ice margin at the time of sampling were extracted from 25 km×25 km resolution passive microwave satellite raster imagery obtained from the Sea Ice Index Version 3.0 product of the National Snow and Ice Data Centre (Fetterer et al., 2017) using a custom script in R.

The foraminiferal assemblage captured in the net is the result of growth over several weeks (Carstens & Wefer, 1992). To a certain degree, the observed composition thus reflects processes acting throughout the habitat traversed by the plankton before being intercepted by the net. To account for the effect of these processes, next to the in-situ parameters, we also analyse two descriptors of the overall oceanographic state of the sampling area (spatial polygon including all the sampling locations present in our compilation) at the time of sampling. These include the average SST of the sampling area derived from the NOAA Optimum Interpolation Sea Surface Temperature V2 (Reynolds et al., 2002) and the average sea-ice extent of the sampling area extracted from the Sea Ice Index Version 3.0 (Fetterer et al., 2017). Data on the projected Arctic sea-ice extent between 2010 and 2090 relative to the month of August under two climate scenarios (RCP 4.5 and RCP 8.5) were obtained from the Climate data store (https://cds.climate.copernicus.eu/). The projected Arctic sea ice coverage is derived from model simulations presented in Khon et al. (2017).

In addition to the physical environment, the foraminiferal population also likely reflects the trophic structure of their habitat. This is often highly correlated with the physical parameters of the environment (sea ice extent, distance from sea-ice edge), but could also act independently. Unfortunately, neither in-situ observations, nor satellite image data are available throughout the sampling period to generate representative and robust estimates of productivity.

### Statistical analyses

We used the obtained dataset to investigate the effect of the environmental parameters and time (i.e., sampling year, since all years the sampling took place in summer) on the composition, total abundance, density, and depth habitat of planktonic foraminiferal species in the Fram Strait. First, we had to rule out the potential influence of the longitudinal gradient of physical properties in the Fram Strait (Fig. 1) on the monitored parameters. Since in most years, the geographical extent of the sampling straddled this gradient, the presence of different hydrographic regimes in the east and in the west Fram Strait (WSC and EGC respectively) could potentially be the dominant factor influencing the planktonic foraminifera community. To test the effect of the longitudinal gradient, we performed a hierarchical cluster analysis using the unweighted pair group method with arithmetic mean (UPGMA) based on the Bray–Curtis dissimilarity index on species density data using the *hclust* function in the package “vegan” (version 2.5-6) (Oksanen et al., 2018) in R and observed the clustering of sample sites assigned to the two hydrographic regimes in the region defined as in Fadeev et al. (2018). This analysis revealed no preferential clustering of samples according to the region (Fig. 1b), indicating that the observed variability is due to factors other than the sampling location.

Changes in the community structure of planktonic foraminifera were then analysed using a multivariate approach. We used nonmetric multidimensional scaling (NMDS) to visualize the similarities of assemblages observed across the stations using the *metaMDS* function from the R package “vegan” (version 2.5-6) (Oksanen et al., 2018). For the NMDS, data by Carstens (1997) were not included in order to eliminate potential biases due to the taxonomic ambiguity in the counts (*N. pachyderma* and *N. incompta* not distinguished, see above). The obtained ordination was used to assess the individual effects of the tested environmental variables on the foraminiferal community by performing BIOENV analysis (Clarke & Ainsworth, 1993). This test allows the identification of variables that best explain the variance in the biological community by calculating a correlation coefficient that is then subjected to a permutation test to determine its significance. Prior to this step, we checked for the presence of collinearity between the environmental variables using the variance inflation factor (VIF) with the *vifstep* function from the R package “usdm” (version 1.1-18). The function calculates the VIF for a set of variables and excludes the highly correlated variables (VIF > 5) (Fenton et al., 2016) from the set through a stepwise procedure. The remaining environmental variables were included in the BIOENV analysis using the *envfit* function from the R package “vegan” with 999 permutations.

Next, generalized linear models (GLM) were applied to assess the effects of time and environmental drivers on the individual density of *N. pachyderma*, *T. quinqueloba* and of the Atlantic species. As we analysed count data, we used the *floor* function in R to derive discrete values from the total concentrations of the three taxonomic groups as a prior step (Zuur et al., 2007) and explored the relationship with the potential predictors with bivariate GLM using the *glm* function in R indicating a quasi-poisson error distribution with log as link function. For the three groups, *N. pachyderma*, *T. quinqueloba* and Atlantic species, the total concentration (dependent variable) was regressed against the sampling year (time), longitude (as a proxy for the two hydrographic regimes), SST, average SST of the sampling area, ice-concentration, distance from ice-margin, AWz, and average sea-ice extent of the sampling area as independent variables. Trait variance explained by individual parameters was calculated using pseudo-R^2^ for Poisson GLMs as 100*(model null deviance-model deviance)/model null deviance (Dobson, 2002). Where more than one predictor displayed a significant effect on the density, VIF was calculated among the variable, variables identified as causing variance inflation were dropped, and the GLMs were re-applied allowing for interactions among the remaining variables.

The relationship between environmental and temporal controllers on the depth habitat (DH) of the resident species *N. pachyderma* and *T. quinqueloba* were investigated through bivariate correlation (Pearson *r*). Square root transformation was performed on *T. quinqueloba* data to obtain symmetric distribution. Multiple linear models were applied to species depth habitat and variables that displayed a significant correlation. The normality of the residuals was checked after the linear model was applied (Zuur et al., 2009). As for the density, in case of more than one predictor displayed a significant correlation with DH, we proceeded to calculate the VIF between the concurrent variables and re-applied the linear model for the remaining variables allowing interactions. Results from models that explained most of the variance (higher pseudo-R^2^ and R^2^) are presented and discussed.

## 5.3 Results

### Species composition

All samples contained an assemblage typical for the polar environment of the Fram Strait, dominated by *N. pachyderma*, which represented 56% of the total assemblage in our compilation. Not all the stations presented the same species proportions: *T. quinqueloba* was the most abundant species in 17 stations. The abundance of the Atlantic species also varied greatly in our compilation ranging from total absence to 27 % of the total assemblage in the samples taken in 2014. The BIOENV analysis revealed that three of the tested variables correlate significantly with the obtained ordination without variance inflation: Year (R^2^ = 0.46, p-value = 0.001), SST of the sampling area (R^2^ = 0.36, p-value = 0.008), and distance from the sea-ice margin (R^2^ = 0.23, p-value = 0.023) (Fig. 3 and Fig. 4b). This indicates that the assemblage composition has changed through time and that at least part of the change can be attributed to changes in the physical environment.

### Species density

The time series of population density reveal a considerable amount of variance within each sampling period, with years with unusually high density for some species, and an apparent trend of increasing density of Atlantic species (Fig. 2a). Potential predictors of the observed trends in population density of the three taxonomic groups were thus investigated using generalised linear model and the results are summarised in Table 1 and in Figure 4. The two best predictors of the density of *N. pachyderma* were longitude (Pseudo-R^2^ = 0.18, p-value = 0.01) and the SST of the sampling area (Pseudo-R^2^ = 0.14, p-value = 0.002), both showing a negative relationship with the concentration of *N. pachyderma* (Fig 4a). The final model including the summing effects of the two predictors explained 38% of the total variance. The SST of the sampling area was also negatively associated with the density of *T. quinqueloba* (Pseudo-R^2^ = 0.14, p-value = 0.002), but the variable was removed due to high collinearity (VIF > 5). The remaining two predictors, year of sampling and sea-ice extent had a VIF < 2 and were included in the final model along with their interactions explaining 51% of the observed variance. Only the year of sampling alone was identified as a significant predictor of the total density of the Atlantic species (Pseudo-R^2^ = 0.17, p-value = 0.02).

**Figure 2.**
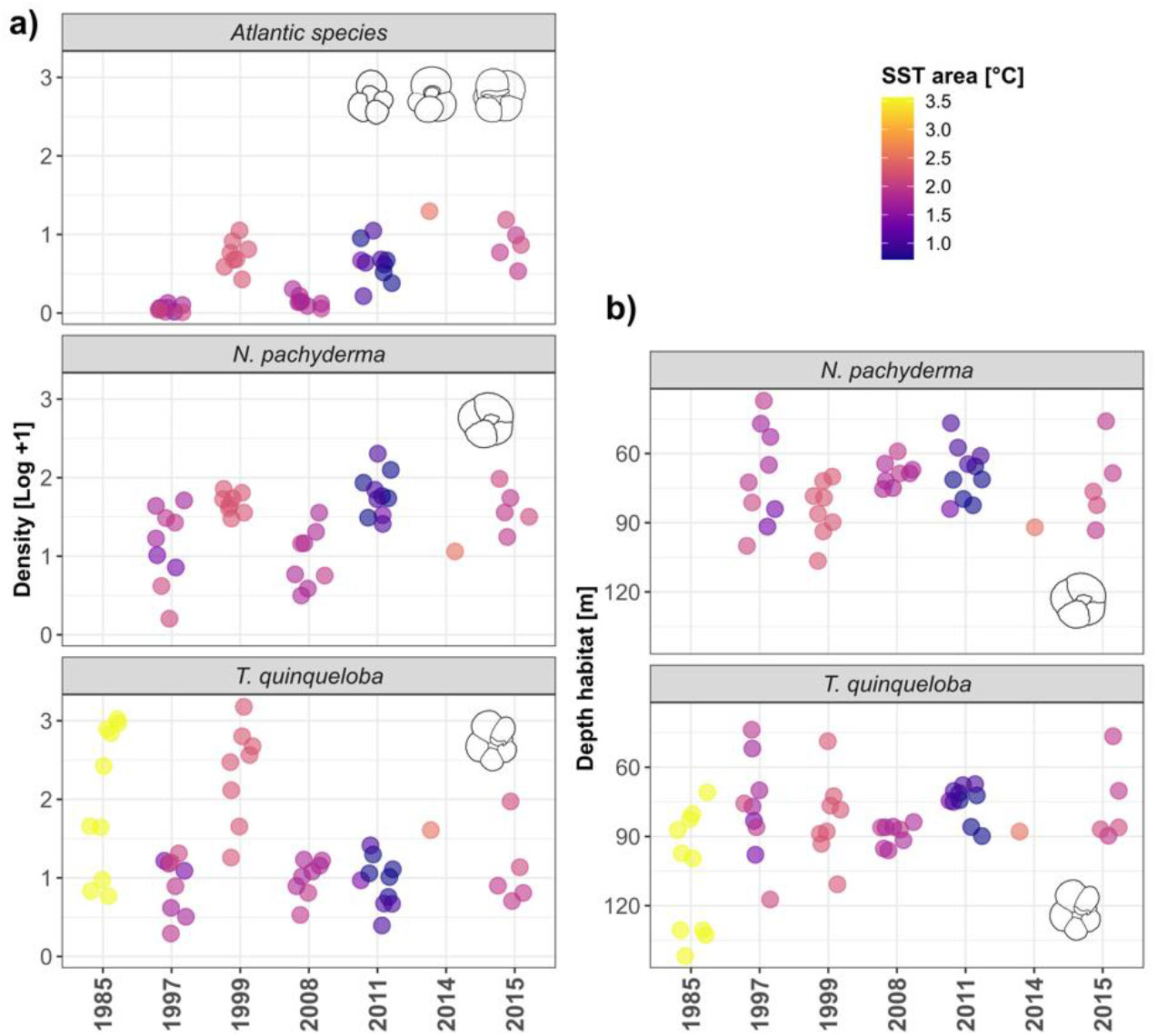
a) Density and Depth habitat (b) of planktonic foraminifera species plotted against year. Colour indicates temperature of the area at the time of sampling.

**Figure 3.**
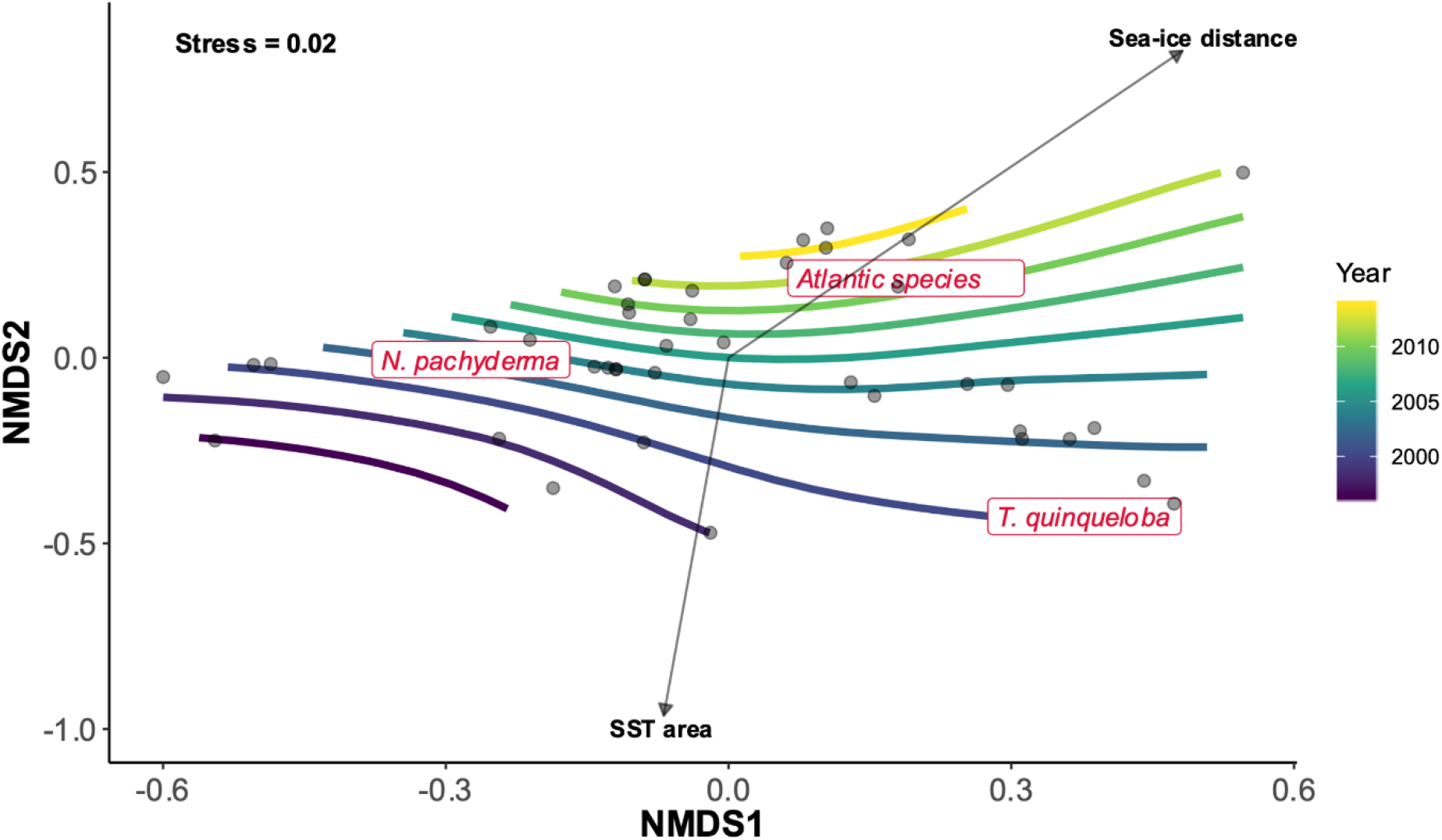
NMDS ordination based on Bray-Curtis similarities Index of planktonic foraminifera abundances with fitted environmental vectors. Contour lines were derived from surface fitting (GAM) of the variable sampling year.

**Figure 4.**
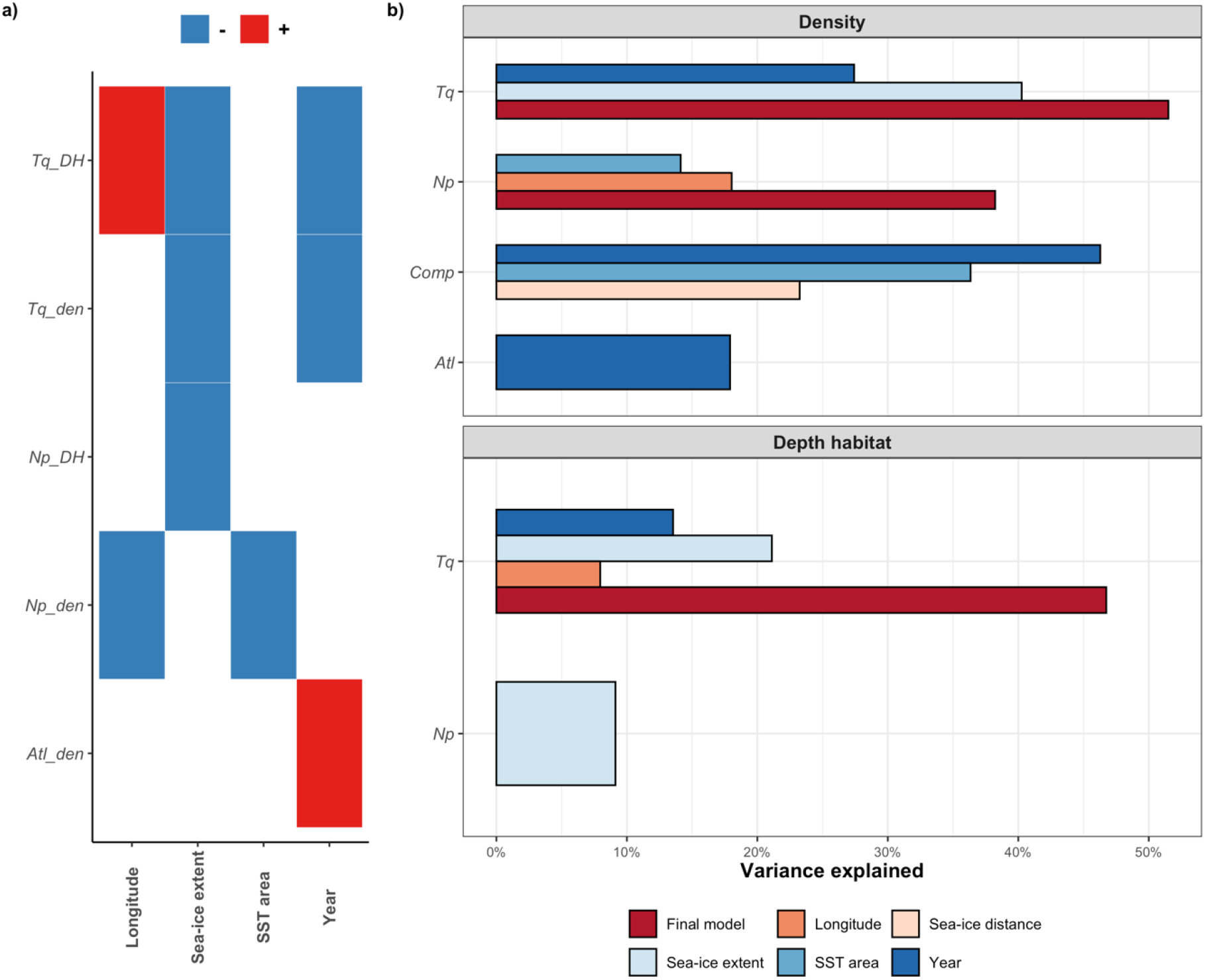
a) Heat-map showing the direction of the relationship with tested environmental variables and modelled responses. b) Bar plot showing amount of variance explained by the singular predictor and the final model. (Abbreviations: Np= *Neogloboquadrina pachyderma*, Tq= *Turborotalita quinqueloba*, Atl= Atlantic species, DH= depth habitat, den= density, Comp= species composition).

**Table 1.** Results of the Generalised linear models and mixed linear models.

|  | *p* - value | Pseudo / Adj. R<sup>2</sup> |
| --- | --- | --- |
| ***N. pachyderma* Density** |  |  |
| Longitude | 0.01 | 18.03 |
| SST area | 0.02 | 14.12 |
| SST area + Longitude |  | 38.23 |
| ***T. quinqueloba* Density** |  |  |
| Year | 0 | 27.42 |
| SST area | 0 | 29.06 |
| Sea-ice extent | 0 | 40.26 |
| Year * Sea-ice extent |  | 51.5 |
| **Atlantic species Density** |  |  |
| Year | 0.02 | 17.91 |
| ***N. pachyderma* Depth habitat** |  |  |
| Sea-ice extent | 0.03 | 9.13 |
| ***T. quinqueloba* Depth habitat** |  |  |
| Longitude | 0.03 | 7.96 |
| Latitude | 0.02 | 8.53 |
| Year | 0 | 13.54 |
| SST | 0.02 | 8.4 |
| SST area | 0 | 19.14 |
| Sea-ice concentration | 0.01 | 12.5 |
| Sea-ice distance | 0 | 13.54 |
| Sea-ice extent | 0 | 21.11 |
| Longitude * Sea-ice extent * Year | 0 | 46.74 |

### Depth habitat

The two species *N. pachyderma* and *T. quinqueloba* displayed a similar vertical distribution in the water column with average living depths 37–140 m and 44–142 m, respectively (Fig.2b). Factors controlling the variability in the observed depth habitat of the two species were investigated by a linear model. This revealed that sea-ice extent in the sampling area alone was the only significant predictor of the depth habitat of *N. pachyderma* (Adj. R^2^ = 0.091, p-value = 0.03). In contrast, all the variables investigated showed a significant correlation with the variability of the depth habitat of *T. quinqueloba* (Table 1) and only the overall SST in the area had to be excluded because of variance inflation. A combined linear model with interactions still identified three variables as significantly and independently affecting the depth habitat of the species. Longitude, sea-ice extent and sampling year explain together 47 % of the observed variance in the depth habitat of *T. quinqueloba*.

## Discussion

The results of the BIOENV analysis indicate a steady rise in the concentration of Atlantic species throughout the observational period (Fig. 3). The same pattern emerged from the GLM (Table 1, Fig. 4) with the year of sampling significantly correlated with population density of the Atlantic species and explaining 18% of the variance in our observations. The declining trend in SST in the Fram Strait suggests that the observed increase in the abundance of Atlantic species in the region cannot be the result of habitat tracking. Since none of the tested environmental factors was a significant predictor of the density of Atlantic species in the region either, their rising abundance must reflect changes in the “source” region in the Nordic Seas from where the species are advected northward with the AW. An increase in density or a change in phenology of these species in the Nordic Seas would result in higher density in the Fram Strait region even without changes in the intensity of AW inflow. Indeed, evidence from moorings has shown that the variability observed in the advection of “Atlantic” copepods in the Fram Strait reflects their phenology and not the intensity of the AW inflow (Basedow et al., 2018). The invoked changes in Atlantic species population dynamics in the “source” region is consistent with the increasing abundance of planktonic foraminifera in the North Atlantic recorded in Continuous Plankton Recorder (CPR) observations (Beaugrand et al., 2013). Importantly, a recent investigation based on satellite-derived altimetry observations showed that an increase in surface velocity of the North Atlantic Current from 1993 to 2016, resulted in a northward shift in the spatial distribution of the coccolithophore *Emiliania huxleyi* (Oziel et al., 2020).

In contrast to the rising abundance of the Atlantic expatriates, the subpolar resident species *T. quinqueloba* shows decreasing population density through time (Figures 2, 3 and 4) leading to lower proportions in the planktonic foraminiferal community (Fig. 3). Contrary to the non-resident, advected Atlantic species, the Fram Strait region is the primary habitat of *T. quinqueloba* (Schiebel et al., 2017) and its abundance in the Fram Strait is not reflecting AW inflow. However, this species is known to prefer warmer, subpolar, waters and is largely absent from the Arctic proper (Carstens et al., 1997; Manno & Pavlov, 2014; Pados & Spielhagen, 2014; Volkmann, 2000). Therefore, the increasing sea-ice export and decreasing SST in the Fram Strait make the region less suitable for this species. Indeed, next to time, a significant component of the variability in the abundance of this species can be explained by local properties in the Fram Strait, in a direction consistent with the above hypothesis: the species is less abundant where/when sea-ice cover is more extensive (Table 1, Fig. 4). Since the local conditions in the Fram Strait are highly variable, a single observation like that by Manno & Pavlov (2014), would easily appear to indicate an opposing trend, highlighting the necessity and merit of the long-term replicated data series presented in this study.

Thus, the changing abundance of *T. quinqueloba* appears to be consistent with habitat tracking, responding to the temporal evolution of local conditions in the Fram Strait. This conclusion is further supported by the observed concomitant shallowing of the vertical habitat of this species through time (Fig. 4), which is reflecting the shallower habitat of the species in the presence of sea ice (Table 1). Previous observations on *T. quinqueloba* in the Fram Strait also showed shallower habitat in the presence of sea ice (Carstens et al., 1997; Volkmann, 2000). The current increase in sea ice in the Fram Strait thus acts to reduce the population density of this species and shoal its vertical habitat, both occurring in the direction consistent with habitat tracking.

Consistently with the increasing sea-ice extent and decreasing temperature, the habitat of the Fram Strait remains suitable for the polar species *N. pachyderma*, which shows no significant temporal trend in its density or vertical habitat (Fig. 2). Instead, the variability in these parameters can be explained by local parameters with higher density and shallower habitat occurring when and where the sea-ice cover is more extensive (Table 1, Fig. 4). Peaks in *N. pachyderma* density in cold polar waters were observed in previous studies (Manno & Pavlov, 2014; Volkmann, 2000) as well as its high occurrence along the sea-ice margin considered, where higher primary production by diatoms represents a major food source for this species (Greco et al., 2021). The habitat shoaling towards sea ice is entirely consistent with a recent analysis of factors affecting the vertical habitat of this species (Greco et al., 2019).

Our in-situ, vertically, resolved observations of three decades of plankton change in the Fram Strait provide direct evidence that trends in population density are associated with significant shifts in the vertical position of the involved species. This observation is significant, as it could not have been derived from CPR or sediment trap devices or remote sensing of the ocean surface. The existence of systematic vertical shifts in plankton populations has significant consequences for biogeochemical cycling in the upper ocean (Bianchi et al., 2013). In addition, the changes in plankton vertical habitat in the Fram Strait may affect species interactions with other resident or immigrant Atlantic species, as vertical niche partitioning among closely related species of zooplankton is an important mechanism of adaptation to the Arctic environment (Kosobokova et al., 2011). In light of these observations, we postulate that the assessment of future changes in the marine biota in the Arctic gateway must also consider the vertical dimension of the pelagic habitat (Gluchowska et al., 2017; Jorda et al., 2020; Knutsen et al., 2017; Kosobokova et al., 2011).

Overall, we thus show that plankton in the Arctic gateway is assuming an unusual composition, with the resident species shifting towards more polar taxa and shallower habitat, tracking local environmental change, being confronted with increasing abundance of Atlantic expatriates, rising due to processes favouring their growth in the Nordic Seas. Since there is no reason to believe that this observation based on planktonic foraminifera should not apply to other plankton groups, this shift in community composition likely alters the diversity of planktonic communities, in turn affecting the established food webs of the involved species (Griffith et al., 2019; Kortsch et al., 2015). At present, the increased sea-ice export in the Fram Strait compensates the overall regional warming in the Arctic, muting the changes in plankton communities in the region. Indeed, in the adjacent Barents Sea, in absence of sea-ice export, the Atlantification of the foraminiferal community appears stronger (Ofstad et al., 2020), likely further enhanced by import of nutrients that promotes phytoplankton production (Lewis et al., 2020). This means that once the ice export in the Fram Strait ceases to be fuelled by the increasing Arctic sea ice reduction (Årthun et al., 2021; Guarino et al., 2020), the planktonic community will likely abruptly shift to a completely different state with more Atlantic and more non-sea-ice species, possibly impacting the carbon export of the region (Anglada-Ortiz et al., 2021). Observational data of sea-ice extent and future predictions plotted in Figure 1d show that, in the Fram Strait, the trend towards an increase in sea-ice export seems to have already reached its maximum. The projections point at a further reduction showing that by the year 2050, the sea-ice extent in the area will attain values below the variability of the observational era in the last four decades (Fig. 1d). Thus, this would be the time when we can expect the regime shift to occur. Acting as the gateway to the Arctic, this rapid shift in the Fram Strait will likely propagate into the Arctic proper.

## Data availability

All new data on which this study is based will be deposited on the public repository PANGAEA. The data as used in this study, including the environmental data and biological data sources, are available on Zenodo (https://doi.org/10.5281/zenodo.5266464).

## Acknowledgements

The masters and crews of the *R/V Polarstern* and *R/V Helmer Hanssen* are gratefully acknowledged for support of the work during the PS93 and HH/14 cruises. Sampling during the PS93.1/ARK-XXIX/2.1 cruise was supported by the Byrd Polar and Climate Research Center, Columbus, Ohio, United States and the National Science Foundation Paleo Perspectives on Climate Change (P2C2) program #1404370. The CAGE 14.4 (HH/14) was supported by the Research Council of Norway through the project Effects of Ocean Chemistry Changes on Planktic Foraminifera in the Fram Strait: Ocean Acidification from Natural to Anthropogenic Changes, (project no. 216538) and the Centre for Arctic Gas Hydrate, Environment and Climate, UiT (project no. 223259). This research has been supported by the Deutsche Forschungsgemeinschaft (DFG) through the International Research Training Group “Processes and impacts of climate change in the North Atlantic Ocean and the Canadian Arctic” (IRTG 1904 ArcTrain).

## Notes

https://doi.org/10.5281/zenodo.5266464

### Competing Interest Statement

The authors have declared no competing interest.

